# AOP: An R Package For Sufficient Causal Analysis in Pathway-based Screening of Drugs and Chemicals for Adversity

**DOI:** 10.1101/029694

**Authors:** Lyle D. Burgoon

## Abstract

**Summary:** How can I quickly find the key events in a pathway that I need to monitor to predict that a/an beneficial/adverse event/outcome will occur? This is a key question when using signaling pathways for drug/chemical screening in pharmacology, toxicology and risk assessment. By identifying these sufficient causal key events, we have fewer events to monitor for a pathway, thereby decreasing assay costs and time, while maximizing the value of the information. I have developed the “aop” package which uses back-door analysis of causal networks to identify these minimal sets of key events that are sufficient for making causal predictions.

**Availability and Implementation:** The source for the aop package is available online at Github at https://github.com/DataSciBurgoon/aop and can be installed using the R devtools package. The aop package runs within the R statistical environment. The package has functions that can take pathways (as directed graphs) formatted as a Cytoscape JSON file as input, or pathways can be represented as directed graphs using the R/Bioconductor “graph” package. The “aop” package has functions that can perform backdoor analysis to identify the minimal set of key events for making causal predictions.

## 1 INTRODUCTION

Pathways provide a mechanism to biologically anchor and provide context for data from high throughput *in vitro* and *in vivo* screening assays, as well as omics studies. When these pathways are associated with a biological effect, they generally provide a causal path showing how perturbations in metabolite/protein/gene activity/expression following chemical exposure leads to a biological effect. Overlaying screening data onto the pathway yields insights as to the potential information flow and the activation/inactivation of the enzymes/proteins in the pathway. This can give us clues as to whether or not a drug/chemical exposure may lead to a biological effect.

However, there are generally more than one pathway that may lead to a biological effect, and these pathways may interact in a myriad of ways. For instance, if there are two parallel pathways that both converge at the same biological effect, there are many possible situations that can arise. Both pathways may be necessary to activate the biological effect. Alternatively, activation of either pathway in the absence of the other may be sufficient to lead to the biological effect. There may also be cross-talk such that one pathway can interfere with the other, and prevent activation/deactivation of the biological effect.

In the field of toxicology, the adverse outcome pathway (AOP), has recently emerged as a concept that allows for efficient modeling of the necessary key events that ultimately lead to an adverse outcome (Ankley *et al.,* 2010). AOPs are causal networks, where activation of a key event parent leads to either the activation or inactivation of a child key event, and so on, ultimately leading to some adverse outcome.

Although it would be ideal to have a high throughput screening assay for every key event in an AOP, it is simply impractical. Financial factors alone render a toxicology screening program focused on measuring every key event across every conceivable AOP impractical. For cost and efficiency reasons, a smaller number of assays that are sufficient to predict an adverse outcome is needed.

Thus, I have developed an R package, called “aop”, which implements Pearl’s backdoor algorithm to identify those one or more key events that are sufficient to be measured and predict whether or not an adverse outcome is likely. The backdoor algorithm uses causal network analysis to identify the sufficient key event nodes. The package also contains functions that can ingest AOPs constructed in Cytoscape exported as Cytoscape JSON files, as well as AOPs constructed as graphs (graphNEL objects) using the Bioconductor “graph” package.

## 2 ALGORITHM

For the sake of this description, consider a biological pathway to be the same as a mathematical graph. A graph is a set of nodes/key events that are connected to each other through edges/lines/arrows. Because this is a causal graph, the edges are directed, such that if node A activates node B, an arrow is drawn from node A to node B. This is generally the same convention used in most biological pathways. AOPs by definition are causal graphs. The “aop” package requires that the input AOP network be a directed acyclic graph.

Pearl’s backdoor algorithm can be used to identify the nodes between a chemical’s entry into the graph/pathway and the adverse outcome that needs to be measured to make the causal argument that the chemical exposure results in the adverse outcome. In the backdoor algorithm, the first step is to introduce an edge between parent nodes that share a child node. Next, the algorithm identifies the shortest path (using the “igraph” package) starting at the adverse outcome and ending at the key event specified by the user where the chemical enters the pathway. The causal node is the second node in the path returned from igraph, where the first node is the adverse outcome. The algorithm then removes the causal node from the pathway, and re-runs the path-finding algorithm to identify the next causal node, and continues this sequence until there are no paths leading from the adverse outcome to the user-specified key event where the chemical enters.

The package has functions to ingest AOPs drawn in Cytoscape as Cytoscape JSON files. The package can also utilize AOPs stored as graphNEL objects (graphNEL is a class from the graph library). The package vignette and examples contains additional details.

## 3 EXAMPLE

Figure 1 shows an AOP for hepatosteatosis, or non-alcoholic fatty liver disease, based on the Reactome Peroxisomal Lipid Metabolism Pathway (http://www.reactome.org/PathwayBrowser/#DIAGRAM=390918) and other articles (Kay *et al.,* 2011; Reddy, 2001). This AOP network shows two parallel tracks that can both result in hepatosteatosis.

**Fig. 1.**
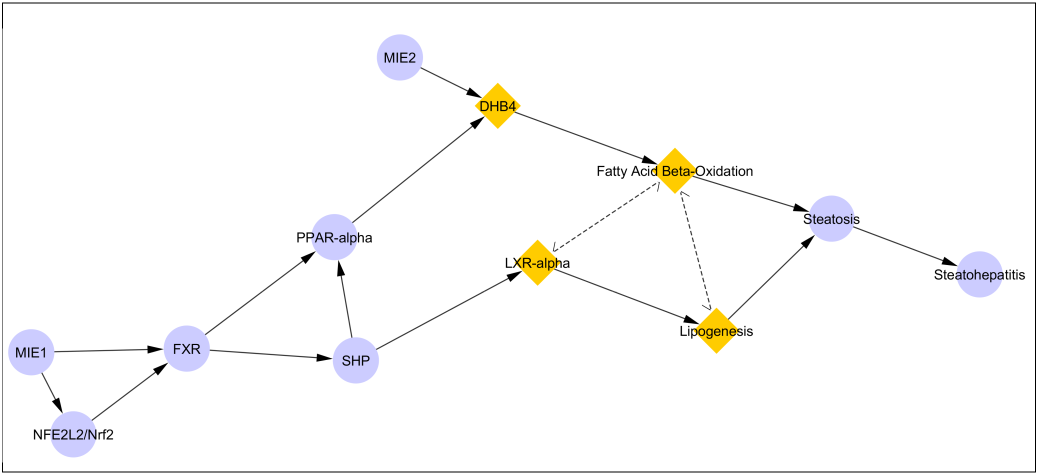
Hepatosteatosis Adverse Outcome Pathway Network. This depicts an AOP network with two parallel paths leading to steatosis and steatohepatitis. The solid arrows are causal linkages between the key events (nodes: circles and diamonds). The nodes MIE1 and MIE2 are 2 hypothesized molecular initiating events where a chemical may perturb the network. The dotted arrows are edges that the algorithm infers. The diamond shaped key events are the sufficient causal ones identified by the algorithm.

The hepatosteatosis adverse outcome pathway network consists of two adverse outcome pathways, both leading to an abnormal accumulation of fat in the liver (hepatosteatosis). Although one could measure all of the key events starting just after the molecular initiating event (MIE; the point where the chemical enters the pathway) through the adverse outcome, this would be costly and time-consuming.

Because adverse outcome pathways are causal networks, the backdoor algorithm provides the capability to identify those key events/nodes that should be monitored to identify causality. Running the backdoor algorithm using MIE2 as the starting point and the steatosis node as the adverse outcome, the algorithm identifies that fatty acid beta-oxidation and lipogenesis both need to be monitored.

If these nodes cannot be monitored due to a lack of assays, the backdoor algorithm can be re-run using fatty acid beta-oxidation or lipogenesis as the ending nodes in the path, keeping MIE2 as the starting point. This identifies that measuring DHB4 and lipogenesis are necessary to inform what is happening at the fatty acid beta-oxidation node. Likewise, measuring LXR-alpha and fatty acid beta-oxidation are necessary to inform us about whether or not lipogenesis is occurring. By combining this information, we can see that if lipogenesis and fatty acid beta-oxidation cannot be measured then we need to measure DHB4 and LXR-alpha to draw conclusions about whether or not steatosis is occurring. By allowing investigators to focus on running assays tied to a smaller number of sufficient key events, organizations can maximize efficiency, decreasing time to decisions and decreasing overall testing costs as compared to having to run assays for all of the key events across the pathway.

## 4 CONCLUSIONS

One key challenge with respect to using pathways to inform decision-making (e.g., go/no-go in drug development based on toxicity, chemical hazard identification) is the identification of the minimal set of key events that need to be monitored through assays.

The backdoor algorithm identifies the nodes that are sufficient for inferring whether an adverse outcome will occur. These key event nodes are the ones that are the most cost-effective (largest value of information) for monitoring, allowing decision-makers to identify if an adverse outcome is likely to occur while minimizing testing costs.

## ACKNOWLEDGEMENTS

I would like to thank John Vandenberg, Debra Walsh, Ila Cote, Stephen Edwards, and Shannon Bell for their comments on a previous draft of this manuscript. I would also like to thank Shannon Bell, Ingrid Druwe, Noffisat Oki, Kyle Painter, and Erin Yost for participating in the code review of the package.

